# Evaluating UV-C sensitivity of *Coxiella burnetii* in Skim Milk using a Bench-Scale Collimated Beam System and Comparative Study with High-Temperature Short-Time Pasteurization

**DOI:** 10.1101/2022.07.25.501439

**Authors:** Brahmaiah Pendyala, Pranav Vashisht, Fur-Chi Chen, Savannah E. Sanchez, Bob Comstock, Anders Omsland, Ankit Patras

**Author notes:** Corresponding Author Ankit Patras, Ph.D., Associate Professor, Food Science & Engineering, Department of Agricultural and Environmental Sciences, College of Agriculture, Human Nutrition and Natural Sciences, Tennessee State University, Nashville TN.

## Abstract

*Coxiella burnetii* is a zoonotic Gram-negative obligate intracellular bacterial pathogen and the causative agent of Query (Q) fever in humans. Contamination of milk by *C. burnetii* as a consequence of livestock infection is a significant public health concern. Effective methods to inactivate *C. burnetii* in milk is a critical aspect of food safety. In this study, we measured optical light attenuation factors; absorption, scattering, and reflection of skim milk (SM) and considered for evaluation of delivered UV dose under stirred conditions. The accuracy of the method followed for the estimation of delivered UV dose in SM was verified by comparative studies of *Escherichia coli* ATCC 25922 inactivation in phosphate buffer (transparent fluid), and humic acid (opaque fluid). Absorption, scattering coefficient, and the reflectance of SM at 254 nm was measured as 19 ± 0.3/cm. 26 ± 0.5/cm and 10.6 %, respectively. SM inoculated with *C. burnetii* was irradiated using a collimated beam device equipped with a low-pressure UV-C_254 nm_ lamp at doses from 0 – 12 mJ·cm^-2^. Results showed a log-linear inactivation of *C. burnetii* in SM with UV-C sensitivity (D_10_) value of 4.1 ± 0.04 mJ·cm^-2^. Similar inactivation kinetics was observed with *Salmonella enterica serovar* Muenchen ATCC BAA 1674 in SM and thereby suggested as a suitable surrogate to *C. burnetii* for pilot scale UV-C processing studies of SM.

## Introduction

*Coxiella burnetii* is a zoonotic Gram-negative obligate intracellular bacterial pathogen and the causative agent of Query (Q) fever in humans. Shedding of *C. burnetii* in the milk and other secretions and excretions of infected cows, goats, and sheep is a significant concern for public health (Enright et al., 1957; Shaw and Voth, 2019; Wittwer et al., 2021). The majority of human infections are asymptomatic or emerge as an acute Q fever - a flu-like illness of differing severity, with symptoms which may include fever, chills, headache, fatigue, malaise, myalgia, arthralgia and a cough. (CFSPH, 2017). In some cases, *C. burnetii* can cause severe syndromes, including reproductive losses and pneumonia, which are also life-threatening in people with pre-existing conditions, including heart valve abnormalities (CFSPH, 2017). During the life cycle, *C. burnetii* transitions between a replicative large cell variant (LCV) and a non-replicative small cell variant (SCV) that accumulates in stationary phase (Coleman et al., 2004). The SCV has an unusual spore-like structure with highly condensed chromatin. *C. burnetii* is highly resistant to environmental factors including heat, making one of the significant human bacterial pathogens in milk (McCaul and Williams, 1981; Codex Alimentarius, 2009; Roest et al., 2013).

Heat inactivation studies of *C. burnetii* indicate that current thermal pasteurization conditions could reduce the *C. burnetii* more effectively than demanded (5 log_10_ reduction) in the Codex Alimentarius (Latin for food code, refers international food code established by United Nations. (Enright et al., 1957; Codex Alimentarius, 2009; Wittwer et al., 2021). However, the temperatures used in thermal pasteurization may significantly reduce milk quality, including alteration in the sensorial and nutritional profile of the product, protein denaturation, undesired changes to milk fat globules; (Garcia-Amezquita et al., 2009; Cappozzo et al., 2015; Gunter-Ward et al., 2018). Also, high operational cost makes thermal pasteurization not feasible for small scale dairy units. Consumers prioritize raw dairy products because of their typical sensorial characteristics and other health benefits such as decreased incidence of asthma, allergies, and respiratory infections, which suggest immune benefits of dairy proteins are lost upon heat processing (Buchin et al.,1998; Braun-Fahrländer and Von Mutius, 2011). To preserve the benefits of raw dairy products without compromising their safety warrants the study of non-thermal technology for the processing of dairy products (Gunter-ward et al., 2018).

UV-C light processing has been considered one of the most promising technologies for pathogen inactivation of milk due to its low energy consumption, the possibility of continual operation, and the fact that it does not generate any chemical byproduct (Patras et al., 2021). In a recent decision from the European Union commission, UV-treated (1045 J/l) milk was approved to be marketed with an extended shelf-life (EFSA Scientific Committee, 2016). Antimicrobial properties of UV-C light at 254 nm have been extensively studied against vegetative bacteria, spore forms, viruses, fungi, algae, and protozoa (Malayeri et al., 2016). Little et al. (1980) studied the effect of UV light on *C. burnetii* in suspensions. They reported the inactivation of *C. burnetii* (10^8^ organisms per mL) at UV treatment conditions of 600 µW/cm^2^ for 15 s at a distance of 10 cm and penetration depth of 1 mm. But the authors did not measure the optical attenuation of test fluid, did not report the average dose, and measured the effect of UV treatment on *C. burnetii* indirectly by detecting serum antibodies in mice. Indeed, direct measurement of *C. burnetii* viability via analysis of colony forming units has only recently become possible (Sanchez et al 2018).

Bolton et al. (2003) reported a standard method for estimating UV-C sensitivity in UV-C absorbing fluids. However, in addition to absorbance, some fluids (e.g., milk) can scatter the UV-C light, which needs to be considered to estimate average delivered dose/fluence and thereby UV-C sensitivity. Therefore, the objectives for this study are to (i) develop method for dose measurement and estimate microbial UV-C sensitivity in SM; (ii) evaluate the UV-C sensitivity of *C. burnetii* in SM; (iii) conduct comparative HTST pasteurization study; and (iv) identify a bacterial surrogate with similar UV-C sensitivity to circumvent slower growth rates of *C. burnetii* for further continuous UV-C systems validation.

## Materials and method

### Bacterial culture conditions and titer enumeration

Axenic culture of a chloramphenicol resistant strain of *C. burnetii* Nine Mile phase II (NMII; RSA 439, clone 4) was propagated in acidified citrate cysteine medium-2 (ACCM-2) as described previously (Omsland et al., 2009; Omsland et al., 2011; Sanchez et al., 2018;). *C. burnetii* was stored at −80° C in ACCM-2 supplemented with 10% DMSO. For enumeration of CFUs, samples were diluted serially in ACCM-2 inorganic salts before plating on solid (final 0.25% w/vol. agarose) ACCM-2 supplemented with 500 μM tryptophan and 1.5 *µ*g.ml^-1^ chloramphenicol. Plates were incubated for 9–10 days in a tri-gas incubator at 37 °C with 5% CO_2_ and O_2_ (Omsland et al., 2011; Sanchez et al., 2018). To correlate the number of viable bacteria as measured by CFU analysis to the total number of bacteria used for inoculation, *C. burnetii* genome equivalents (GE) were quantified via detection of the *C. burnetii* gene CBU1206 using a CFX96 Real-Time PCR Detection System (Bio-Rad Laboratories, Hercules, CA) (Beare et al. 2012; Sanchez et al., 2018). *C. burnetii* GEs were extrapolated from a standard curve prepared using recombinant CBU1206. *Escherichia coli* ATCC 25922 and *Salmonella* enterica serovar Muenchen ATCC BAA 1674 were obtained from American Type Culture Collection (ATCC) and propagated in Tryptic soy broth (TSB) and harvested as reported earlier (Pendyala et al., 2021; Vashisht et al., 2021). To enumerate the CFU titer, appropriate dilutions in peptone water (in 0.1% PW) were plated in duplicate onto Tryptic soy agar (Oxoid Ltd., Basingstoke, UK) plates and incubated for 24 hours at 37 °C.

### Preparation of test fluid microbial suspensions

Ultra-high temperature (UHT) processed SM from Parmalat Canada, Humic acid (adjusted pH to 7.0) from Agricultural Services of America, Inc. in Lake Panasofskee, Florida, USA, and phosphate buffer saline (PBS, pH 7.0) were used as test fluids. To remove ACCM-2 and DMSO from the test samples, *C. burnetii* stocks were washed twice with 0.1% (w/v) phosphate-buffered saline (PBS) (Becton Dickinson, New Jersey, US) using centrifugation at 3000 × g, 15 min. The test samples were prepared by spiking SM with microorganisms (*C. burnetii, E. coli* or *S*. Muenchen) at concentration > 10^8^ CFU.mL^-1^ for UV-C irradiation and HTST exposures.

### Measurement of optical properties

Optical properties of SM inoculated with bacteria were measured using a double beam Cary 100 Spectrophotometer (Varian, USA) equipped with a 6-inch single Integrating Sphere (Labsphere, DRA-CA-30, USA) to estimate scattered light at 254 nm wavelength (Shenoy and Pal, 2008). Thin quartz cuvettes (0.08 mm path-length) were prepared and used to measure optical properties. The transmittance and reflectance (diffuse reflectance) of light were collected by the integrating sphere when the sample was placed at the entrance and exit ports, respectively (Gunter-ward et al., 2018). The amount of light transmitted and reflected by the quartz cuvettes was also quantified and considered to estimate absorption, scattering coefficients, and reflectance. The refractive index (RI) of SM samples in the range of 365 nm to 706 nm was measured at 20 ºC using the digital refractometer (Schmidt + Haensch GmbH & Co, Germany). Then a 5^th^-order polynomial fit was used to extrapolate the refractive index at 254 nm. The inverse adding doubling (IAD) program was used to compute absorption and scattering coefficients by applying total transmittance, reflectance, and refractive index as input values (Prahl et al., 1999). Ultraviolet transmittance (UVT-%/cm), which indicates the fraction of incident light transmitted through a material over a 1 cm path, was calculated as per equation 1.

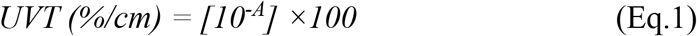

where A represents the absorbance (base_10_) of the test fluid at 254 nm for a 1cm path.

### UV system, dose calculation, and test fluid treatment

A custom-made collimated beam apparatus (Pendyala et al., 2019) was used to irradiate test fluids (Figure 1). The system design followed the recommendations of Bolton and Linden (2003) and utilized a low-pressure mercury vapor arc lamp emitting primarily at 253.7 nm. Irradiance at the position of the fluid surface was measured using calibrated radiometer ILT1700 with SED240 detectors, each equipped with a quartz W diffuser and NS254 spectral filter to ensure that only 254 nm radiation was measured (International Light Technologies, Peabody, MA, USA). The delivered UV dose (fluence) was calculated as the product of the volume average of the fluence rate in the sample and the exposure time by assuming perfect mixing of the sample by the stir bar. 2 mL of test fluid (SM or humic acid or PBS) microbial suspensions treated in 10 mL beaker (Optical path length of 6 mm) at UV dose ranges from 0 – 17 mJ/cm^2^ (n = 3). Since SM scatters UV-C light, scattering factor (*S*) was estimated by radiative transport equation (Eq. 2) using computational fluid dynamics (CFD).

**Figure 1.**
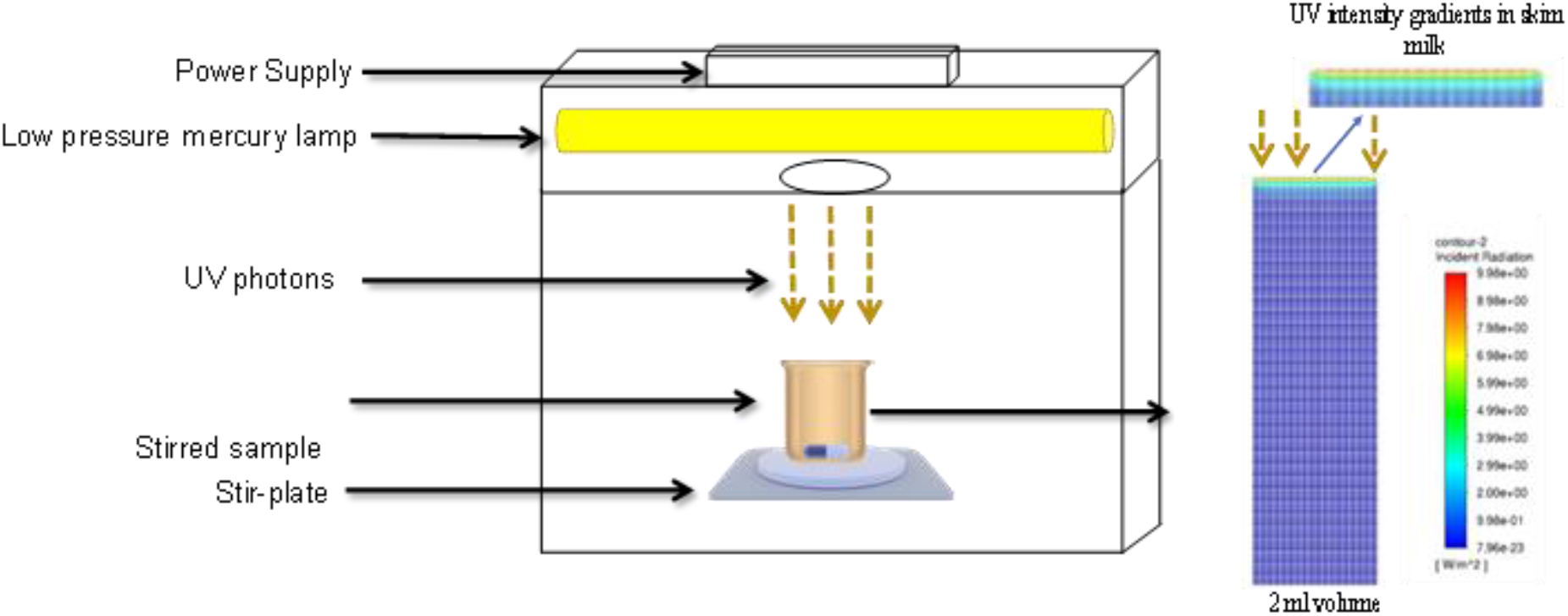
Collimated beam system and computational fluid dynamics analysis (radiation profile) conducted using Fluent program

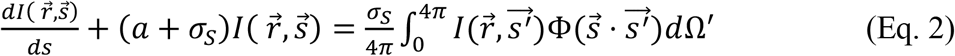

Where *I* is the radiation intensity, 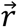 is the position vector, 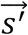 is the scattering direction vector, *s* is the path length, *a* is the absorption coefficient, *n* is the refractive index, *σ*_*S*_ is the scattering coefficient, *σ* is the Stephen-Boltzmann constant, *Φ* is the phase function and Ω^′^ is the solid angle.

Then the average UV fluence rate (*E′avg)* and thereby delivered UV dose *(D)* in the stirred sample was calculated by Eq. 3 (Pendyala et al., 2022)

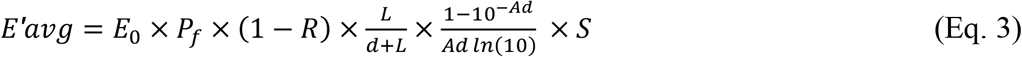

*E*_0_ is the radiometer meter incident irradiance reading at the center top surface of the water in the beaker; *P*_f_ is petri factor; 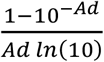 is water factor (1 − *R*) is reflectance factor; and 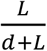 is light divergence factor

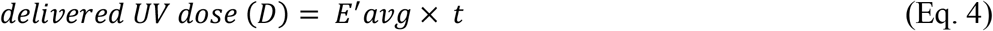

*t* is the exposure time in sec.

### Heat Challenge Studies

The heat challenge studies were conducted using a lab-scale HTST pasteurizer (Kontopodi et al., 2022). The pasteurizer consists of 3 main sections, as shown in Figure 2 a pre-heating section (77 ºC), a holding section (72 ºC for 15 sec), and a cooling unit. The temperature was monitored using a probe. SM inoculated with *C. burnetii* was directly fed into the system using a peristaltic pump (Watson Marlow). The pump was calibrated before the heat challenge studies. Cleaning-in-place (CIP) was conducted before any heat challenge studies. SM was pumped through the heating section, where it achieved the pasteurization temperature of 72 ºC at flowrate of 90 mL.min^-1^ at a 15-sec holding time. At the end of the holding section, SM passed through a cooling coil. In this phase, SM was cooled to a final temperature of 4 ºC±1. 10 ml of samples were collected for enumeration and platting.

**Figure 2.**
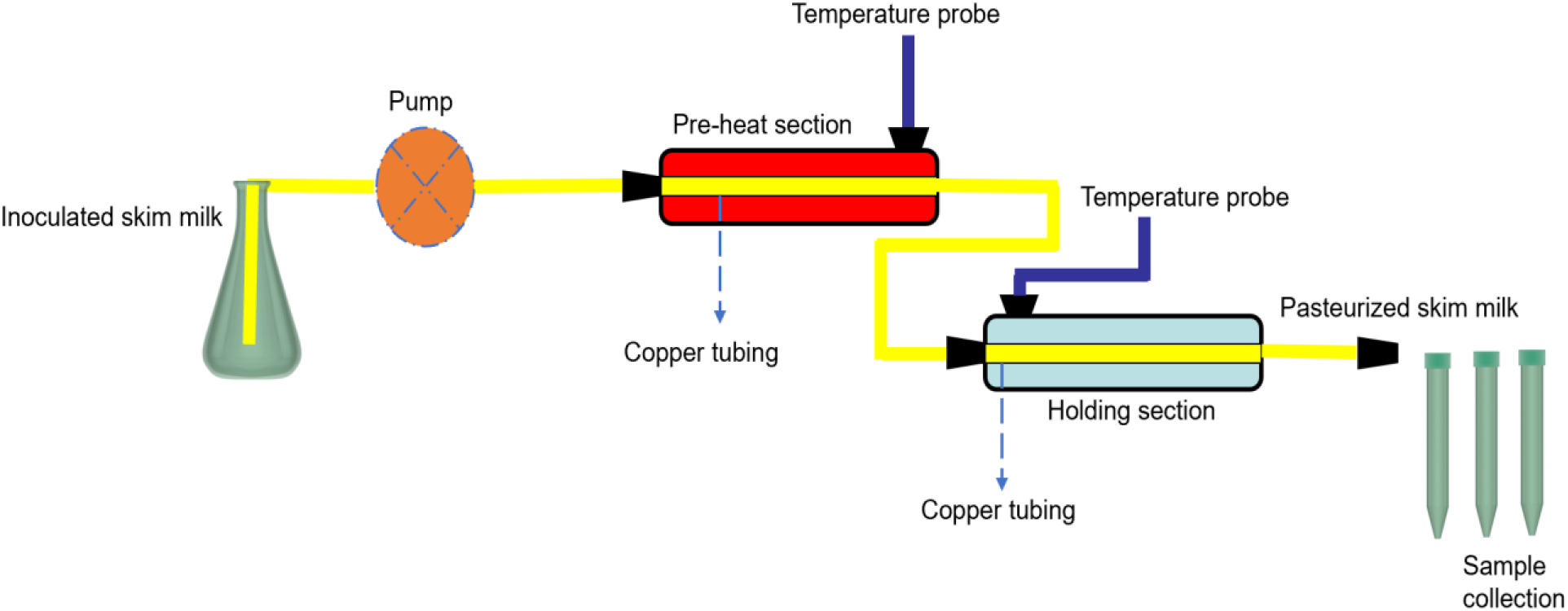
Schematic picture of a lab-built pasteurizer

### Data analysis and Statistics

To assess the inactivation of *E. coli* or *C. burnetii*, the log-linear model available in the GInaFiT tool (a freeware add-in for Microsoft Excel) (Geeraerd et al., 2005) was used to fit the experimental data, and the goodness of fit parameters including R^2^, root mean square error, and rate constants were evaluated. Inactivation kinetics was expressed as follows.

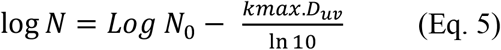

where N and N_0_ are the final and the initial cell number, respectively; *kmax* is the inactivation rate at the highest dose; UV Dose (D_uv_) and D is the decimal reduction value expressed as D_10_ (10% survival in microbial population expressed as mJ·cm^-2^).

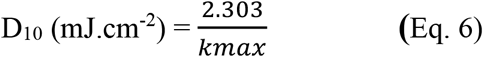

A balanced design with 4 replicates randomized in experimental order was performed for each treatment. Data were reported as means ± one standard deviation from the mean and significance level set to 0.05 (5%).

## Results and discussion

### Optical properties, average fluence rate estimation

Optical properties data indicate that SM strongly absorbs and scatters UV-C light (Table 1). From the measured optical data, it is apparent that the UV-C light has minimal transmission through SM due to aromatic amino acids and other UV-C absorbing organic solutes (Gunter et al., 2018). Light scattering by casein micelles and fat globules causes milk to appear turbid and opaque. The average fluence rate or incident irradiance through the test fluid suspension was 0.0153 mW.cm^-2^ which is estimated by substituting the experimental Petri factor, reflection factor, water factor, divergence factor, and scattering factor in eq.3 (Table 1). Verification of simulated scattering factor and UV dose distribution in SM

**Table 1.** Optical properties and UV doses for skim milk exposure.

| Parameters | Value |
| --- | --- |
| Refractive index | 1.40 ± 0.005 |
| *Absorption coefficient (1/mm) | 4.41 ± 0.03 |
| *Scattering coefficient (1/mm) | 6.02 ± 0.21 |
| Reflection (%) | 10.6 |
| UVT (%/cm) | 6.76E-18 ± 2.05E-18 |
| Surface Irradiance (mW.cm<sup>-2</sup>) | 0.48 ± 0.01 |
| Petri factor | 1.0 |
| Reflection factor | 0.894 |
| Water factor | 0.038 |
| Divergence factor | 0.968 |
| Scattering factor | 0.92 |
| Average fluence rate (mW.cm<sup>-2</sup>) | 0.0153 |

\*Coefficients expressed as Base-e, values reported as mean ± standard deviation

To verify the estimated scattering factor from CFD, comparative microbial (*E. coli*) inactivation studies in SM and humic acid with the same UV-C absorbance as SM without scattering were conducted. The data show there is no significant difference in microbial inactivation kinetics in both SM and diluted humic acid (Figure 3), confirms the accuracy of calculated scattering factor. Dose distribution throughout the fluid domain is a crucial parameter to estimate UV-C sensitivity of microorganisms. Efficient dose distribution conditions provide uniform dose to all microbial particles and thereby results in log-linear inactivation kinetics in mono microbial populations. To check UV dose distribution in SM under standard experimental stir bar mixing conditions, comparative study with a high UV-C transparent fluid phosphate-buffered saline (PBS) – was conducted at same experimental UV-C doses. The experimental data show there is no significant difference (p > 0.05) between the microbial inactivation kinetics with D_10_ values ranging from 3.20 – 3.32 mJ/cm^2^ and shown log-linear inactivation kinetics in the three different test fluids (Table 2). Therefore, these results indicated that the experimental mixing conditions distributed UV-C dose efficiently in opaque test fluids.

**Table 2.** Exposure times to achieve target UV dose for inactivation of *E. coli* for UV dose validation.

| Target UV dose<br>(mJ.cm<sup>-2</sup>) | Exposure time (sec) |  |  |
| --- | --- | --- | --- |
|  | Phosphate<br>buffer | Humic<br>acid | Skim<br>milk |
| 0 | 0 | 0 | 0 |
| 3 | 32 | 151 | 195 |
| 6 | 64 | 303 | 390 |
| D<sub>10</sub> value (mJ.cm<sup>-2</sup>) | 3.32 ± 0.02 | 3.20 ± 0.10 | 3.31 ± 0.07 |

Note: For PBS – distance from UV lamp for = 30.55 cm; average Fluence Rate (mW.cm<sup>-2</sup>) = 0.093230. For humic acid – distance from lamp for = 18.05 cm; average Fluence Rate(mW.cm<sup>-2</sup>) = 0.019824. For skim milk – distance from lamp for = 18.05 cm; average Fluence Rate (mW.cm<sup>-2</sup>) = 0.015394

**Figure 3.**
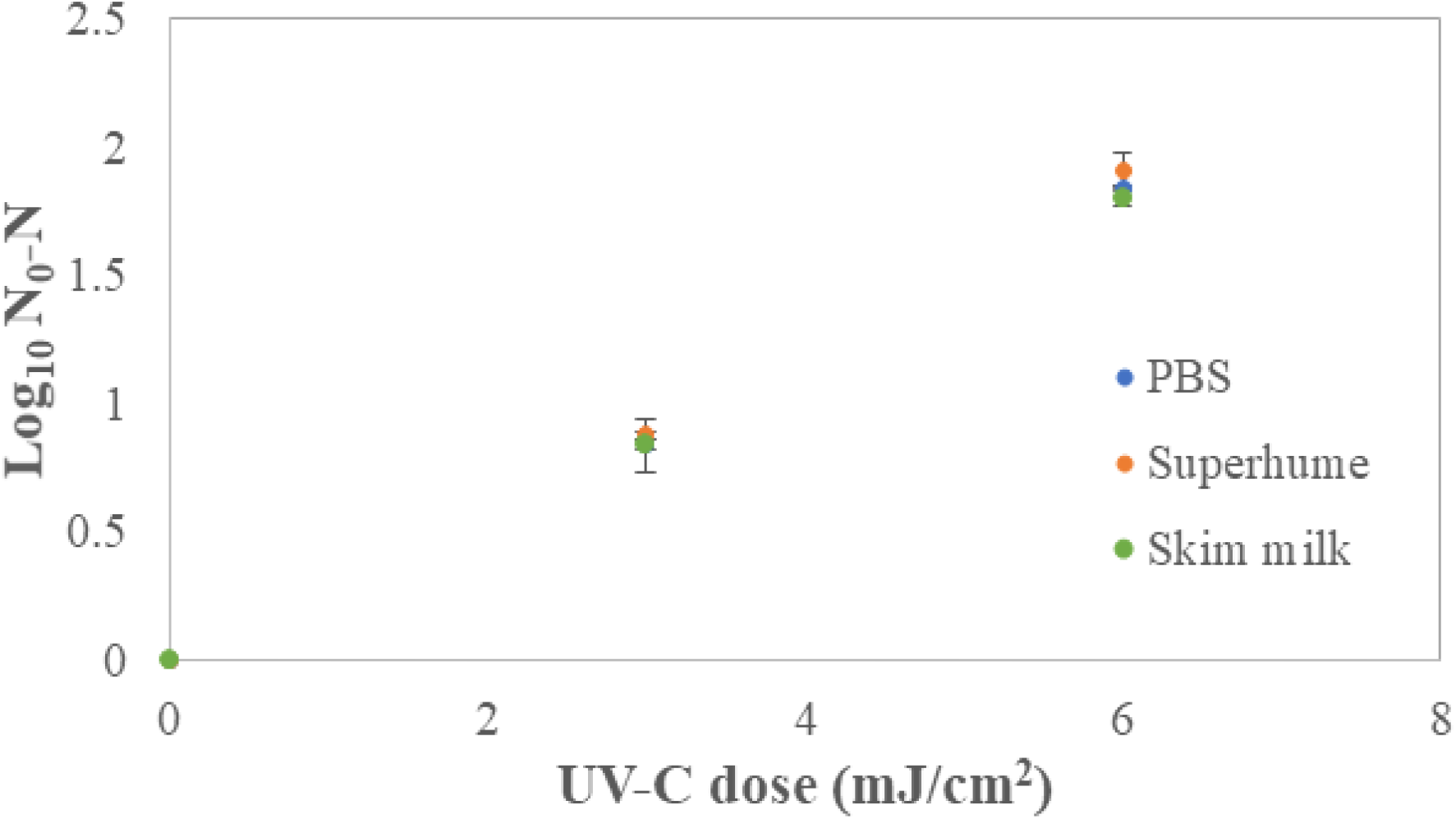
Verification of collimated beam dose delivery using *E. coli* in skim milk, PBS and humic acid. Triplicate irradiations were performed for each dose; all replicates shown on plot, and values shown are averages of duplicate plating of each irradiated sample. Error bars represent range of data

### UV inactivation of *C. burnetii* and identification of bacterial surrogate for validation studies

SM inoculated with *C. burnetii* at approximately 10^8^ CFU.mL^-1^ was exposed to known UV doses of 0, 3, 6, 12 mJ·cm^-2^ in a stirred collimated beam UV system. Figure 4 show *C. burnetii* colonies on ACCM-2 agar plate. The data revealed that > 3 log reduction of *C. burnetii* at a maximum dose of 12 mJ·cm^-2^ (Figure 5). The inactivation kinetics of *C. burnetii* were fitted to a log-linear model with a low (0.35) root mean squared error (RMSE) and higher (0.92) R^2^ (Table 3). The kinetic constant was UV-C sensitivity (D_10_) of *C. burnetii* in SM was estimated as 4.1 ± 0.04 mJ·cm^-2^ with a *k*_*max*_ value of 0.56 cm^2^ /mJ (Table 3). Little et al. (1980) demonstrated the inactivation of *C. burnetii* inactivation in suspension and guinea pig peritoneal macrophages by UV-C light. The authors reported that *C. burnetii* was inactivated at a UV irradiance of 600 μW/cm^2^ for 15 sec at a distance of 10 cm in suspension and macrophages. According to our presented data on the D_10_ value of *C. burnetii*, the performance criterion of 5 log reductions as demanded by the Codex Alimentarius can be achieved at a UV-C dose of 20.5 mJ·cm^-2^.

**Table 3.** Model fitting, Goodness of fit and Predicted UV Dosage for *C. burnetii* in skim milk.

| Parameters | Values |
| --- | --- |
| kmax (cm<sup>2</sup>.mJ<sup>-1</sup>) | 0.56 |
| R<sup>2</sup> | 0.92 |
| RMSE | 0.36 |
| D<sub>10</sub> (mJ.cm<sup>-2</sup>) | 4.1 |
| Dose required to achieve<br>5 log reduction (mJ.cm<sup>-2</sup>) | 20.5 |

**Figure 4.**
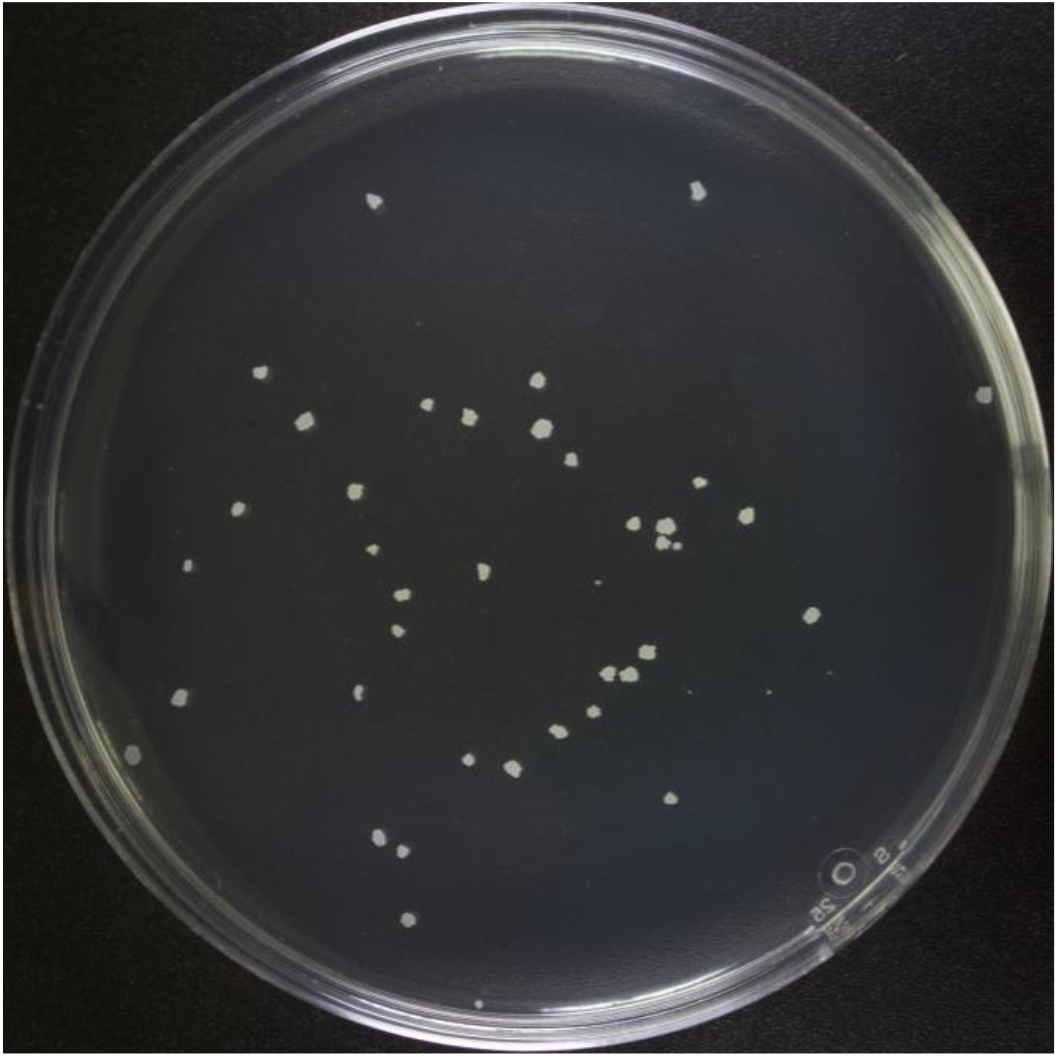
High resolution image of *C. burnetii* growth on a plate

**Figure 5.**
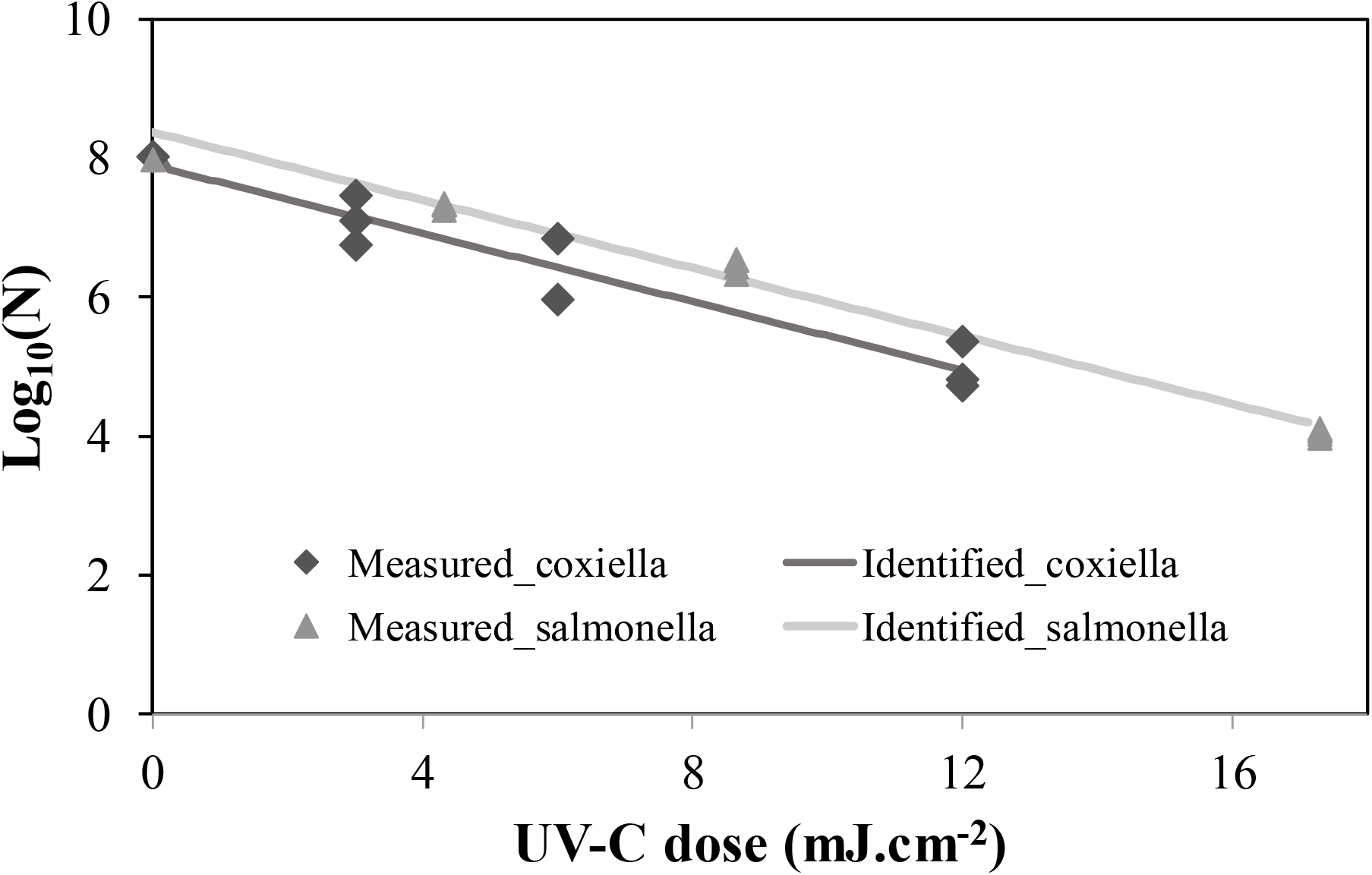
UV-C Inactivation of *C. burnetii* and *Salmonella* enterica serovar Muenchen in SM Triplicate irradiations were performed for each dose; all replicates shown on plot, and values shown are averages of duplicate plating of each irradiated sample. Error bars represent range of data.

While inactivation of *C. burnetii* is a key consideration for milk processing, cultivation of *C. burnetii* requires specialized equipment and expertise. Therefore, to facilitate validation of pilot-scale UV-C systems for inactivation of bacterial pathogens in milk, we sought to identify a bacterial surrogate for *C. burnetii*. Based on the D_10_ of *C. burnetii* (4.1 ± 0.04 mJ·cm^-2^), *Salmonella* strain Muenchen ATCC BAA 1674 with reported D_10_ ranging from 3.9 – 4.3 mJ·cm^-2^ in phosphate-bufferd saline (Gopisetty et al., 2019; Pendyala et al., 2021, Vashisht et al., 2021) was selected and conducted UV-C inactivation studies with these organisms in SM (Figure 5). The results show log linear inactivation kinetics with RMSE 0.1913 and R^2^ > 0.98. Interestingly, the data show *K*_*max*_ (0.56) and D_10_ value (4.1 mJ·cm^-2^) similar to *C. burnetii*, indicative of its suitability as a surrogate for *C. burnetii* for UV-C inactivation studies in SM.

### Comparative HTST pasteurization using SM inoculated with *C. burnetii*

The physical properties of SM and heat exchanger parameters are shown in Table 4. Starting with a *C. burnetii* titer of 3.74×10^8^ CFU.ml^-1^, the heat-dependent log reduction during a holding time of 15 sec at 72 ºC depicted 8 log_10_ reduction of *C. burnetii* in SM, with a D value of approximately 1.8 sec (Table 5). A recent study by Wittwer et al. (2022) on *Coxiella* isolates M, WDK299, and WDK1188 indicated that, in general, all isolates were more susceptible to heat over a temperature range from 60 to 65 ºC with holding times from 15 to 25 sec. For the highest temperature of 65 ºC, the D value was reported as 5.1 to 7.6 sec and predicted reduction of approximately 10.5 log at 72.4 ºC for 15 sec.

**Table 4.** Physical properties of skim milk and heat exchanger parameters.

| Parameters | Value |
| --- | --- |
| Density (kg.m<sup>-3</sup>) | 1.03 |
| Specific heat (J.g<sup>-1</sup> K<sup>-1</sup>) | 3.97 |
| Flowrate (ml.min<sup>-1</sup>) | 90 |
| Rate of heat transfer, Q (J.s) | 404.93 |
| Thermal conductivity, k (J.s<sup>-1</sup> .m<sup>-2</sup>. K<sup>-1</sup>) | 1361.25 |
| Heat exchanger coefficient ($\lambda$) | 1.28E-06 |
| Diameter of the tube (m) | 0.004 |
| Length of tube (m) | 0.54 |
| Area of the tube (m<sup>2</sup>) | 0.0068 |
| Temperature of pre-heat section (°C) | 77 |
| Temperature of heating section (°C) | 72 |
| Holding time (s) | 15 |
| Temperature Cooling section (°C) | 5 (±2) |

**Table 5.** Heat resistance of *C. burnetii* in skim milk.

| **Test condition** | **CFU mL<sup>-1</sup>** | **GE mL<sup>-1</sup>** |
| --- | --- | --- |
| No-treatment control (+) | 3.74x10<sup>8</sup> | 5.38x10<sup>8</sup> |
| No-treatment control (+) | 3.59x10<sup>8</sup> | 4.92x10<sup>8</sup> |
| No-treatment control (+) | 3.32x10<sup>8</sup> | 5.67x10<sup>8</sup> |
| 72 °C / 15 sec | 0.0 (not detected) | 5.20x10<sup>8</sup> |
| 72 °C / 15 sec | 0.0 (not detected) | 5.20x10<sup>8</sup> |
| 72 °C / 15 sec | 0.0 (not detected) | 5.20x10<sup>8</sup> |

### Conclusions

In conclusion, methods to determine optical attenuation properties of UV-C light and eventually calculate the average fluence rate in highly opaque scattering fluids (i.e., SM) were developed and verified the accuracy with experimental results. Importantly, UV-C sensitivity of *C. burnetii* in SM was demonstrated and reported D_10_ value of 4.1± 0.04 mJ·cm^-2^ and predicted UV-C dose of 20.5 mJ·cm^-2^ for 5 log reduction of *C. burnetii* in SM (requirement as per Codex Alimentarius). We have also identified *S*. Muenchen as a suitable surrogate to *C. burnetii* for UV-C treatment validation studies of SM. In comparison, the HTST pasteurization study showed > 8 log reduction of *C. burnetii* in SM. The presented data is critical for the development of non-thermal UV-C pasteurization systems for the processing of SM.

## Acknowledgement

We thank Drs. Bob Heinzen and Paul Beare, Rocky Mountain Laboratories, NIAID, NIH, for sharing the chloramphenicol resistant mutant of *C. burnetii* used in this project.

## References

Malayeri, A. H., Mohseni, M., Cairns, B., Bolton, J. R., Chevrefils, G., Caron, E., … & Linden, K. G. (2016). Fluence (UV dose) required to achieve incremental log inactivation of bacteria, protozoa, viruses and algae. IUVA News, 18(3), 4–6.

Enright, J. B., Sadler, W. W., & Thomas, R. C. (1957). Thermal inactivation of Coxiella burnetii and its relation to pasteurization of milk (No. 517). US Department of Health, Education, and Welfare, Public Health Service.

Sanchez, S. E., Vallejo-Esquerra, E., & Omsland, A. (2018). Use of axenic culture tools to study Coxiella burnetii. Current Protocols in Microbiology, 50(1), e52.

Beare, P. A., Larson, C. L., Gilk, S. D., & Heinzen, R. A. (2012). Two systems for targeted gene deletion in Coxiella burnetii. Applied and environmental microbiology, 78(13), 4580–4589.

Bolton, J. R., & Linden, K. G. (2003). Standardization of methods for fluence (UV dose) determination in bench-scale UV experiments. Journal of environmental engineering, 129(3), 209–215.

Braun-Fahrländer, C., & Von Mutius, E. (2011). Can farm milk consumption prevent allergic diseases?. Clinical & Experimental Allergy, 41(1), 29–35.

Buchin, S., Delague, V., Duboz, G., Berdague, J. L., Beuvier, E., Pochet, S., & Grappin, R. (1998). Influence of pasteurization and fat composition of milk on the volatile compounds and flavor characteristics of a semi-hard cheese. Journal of Dairy Science, 81(12), 3097–3108.

Cappozzo, J. C., Koutchma, T., & Barnes, G. (2015). Chemical characterization of milk after treatment with thermal (HTST and UHT) and nonthermal (turbulent flow ultraviolet) processing technologies. Journal of Dairy Science, 98(8), 5068–5079.

Codex Alimentarius (2009). Code of Hygienic Practice for Milk and Milk Products. CAC/RCP 57–2004, 2nd Edn. Rome: Codex Alimentarius Commission.

Coleman, S. A., Fischer, E. R., Howe, D., Mead, D. J., & Heinzen, R. A. (2004). Temporal analysis of Coxiella burnetii morphological differentiation. Journal of bacteriology, 186(21), 7344–7352.

EFSA Panel on Dietetic Products, Nutrition and Allergies (NDA). (2016). Safety of UV-treated milk as a novel food pursuant to Regulation (EC) No 258/97. EFSA Journal, 14(1), 4370.

Enright, J. B., Sadler, W. W., & Thomas, R. C. (1957). Pasteurization of milk containing the organism of Q fever. American journal of public health and the nations health, 47(6), 695–700.

Garcia-Amezquita, L. E., Primo-Mora, A. R., Barbosa-Cánovas, G. V., & Sepulveda, D. R. (2009). Effect of nonthermal technologies on the native size distribution of fat globules in bovine cheese-making milk. Innovative Food Science & Emerging Technologies, 10(4), 491–494.

Geeraerd, A. H., Valdramidis, V. P., & Van Impe, J. F. (2005). GInaFiT, a freeware tool to assess non-log-linear microbial survivor curves. International journal of food microbiology, 102(1), 95–105.

Gopisetty, V. V. S., Patras, A., Pendyala, B., Kilonzo-Nthenge, A., Ravi, R., Pokharel, B., … & Sasges, M. (2019). UV-C irradiation as an alternative treatment technique: Study of its effect on microbial inactivation, cytotoxicity, and sensory properties in cranberry-flavored water. Innovative food science & emerging technologies, 52, 66–74.

Gunter-Ward, D. M., Patras, A., S. Bhullar M., Kilonzo-Nthenge, A., Pokharel, B., & Sasges, M. (2018). Efficacy of ultraviolet (UV-C) light in reducing foodborne pathogens and model viruses in skim milk. Journal of Food Processing and Preservation, 42(2), e13485.

Kontopodi, E., Boeren, S., Stahl, B., van Goudoever, J. B., Van Elburg, R. M., & Hettinga, K. (2022). High-Temperature Short-Time Preserves Human Milk’s Bioactive Proteins and Their Function Better Than Pasteurization Techniques With Long Processing Times. Frontiers in pediatrics, 9, 798609.

Little, J. S., Kishimoto, R. A., & Canonico, P. G. (1980). In vitro studies of interaction of rickettsia and macrophages: Effect of ultraviolet light on Coxiella burnetii inactivation and macrophage enzymes. Infection and Immunity, 27(3), 837–841.

McCAUL, T. F., & Williams, J. C. (1981). Developmental cycle of Coxiella burnetii: structure and morphogenesis of vegetative and sporogenic differentiations. Journal of bacteriology, 147(3), 1063–1076.

Omsland, A., Beare, P. A., Hill, J., Cockrell, D. C., Howe, D., Hansen, B., … & Heinzen, R. A. (2011). Isolation from animal tissue and genetic transformation of Coxiella burnetii are facilitated by an improved axenic growth medium. Applied and environmental microbiology, 77(11), 3720–3725.

Omsland, A., Cockrell, D. C., Howe, D., Fischer, E. R., Virtaneva, K., Sturdevant, D. E., … & Heinzen, R. A. (2009). Host cell-free growth of the Q fever bacterium Coxiella burnetii. Proceedings of the National Academy of Sciences, 106(11), 4430–4434.

Patras, A., Bhullar, M. S., Pendyala, B., & Crapulli, F. (2021). Ultraviolet treatment of opaque liquid foods: from theory to practice.

Pendyala, B., Patras, A., Gopisetty, V. V. S., & Sasges, M. (2021). UV-C inactivation of microorganisms in a highly opaque model fluid using a pilot scale ultra-thin film annular reactor: Validation of delivered dose. Journal of Food Engineering, 294, 110403.

Pendyala, B., Patras, A., Gopisetty, V. V. S., Sasges, M., & Balamurugan, S. (2019). Inactivation of Bacillus and Clostridium spores in coconut water by ultraviolet light. Foodborne pathogens and disease, 16(10), 704–711.

Pendyala, B., Patras, A., Gopisetty, V. V. S., Vashisht, P., & Ravi, R. (2022). Inactivation of B. cereus Spores in Whole Milk and Almond Milk by Novel Serpentine Path Coiled Tube UV-C System. bioRxiv.

Prahl, S. (1999). Optical property measurements using the inverse adding-doubling program. Oregon Medical Laser Center, St. Vincent Hospital, 9205.

Roest, H. I., Bossers, A., van Zijderveld, F. G., & Rebel, J. M. (2013). Clinical microbiology of Coxiella burnetii and relevant aspects for the diagnosis and control of the zoonotic disease Q fever. Veterinary Quarterly, 33(3), 148–160.

Shaw, E. I., & Voth, D. E. (2019). Coxiella burnetii: a pathogenic intracellular acidophile. Microbiology, 165(1), 1.

The Center for Food Security and Public Health (CFSPH). (2017). Q Fever [Fact sheet]. https://www.cfsph.iastate.edu/Factsheets/pdfs/q_fever.pdf

Vashisht, P., Pendyala, B., Gopisetty, V. V. S., & Patras, A. (2021). Modeling and validation of delivered fluence of a continuous Dean flow pilot scale UV system: monitoring fluence by biodosimetry approach. Food Research International, 148, 110625.

Wittwer, M., Hammer, P., Runge, M., Valentin-Weigand, P., Neubauer, H., Henning, K., & Mertens-Scholz, K. (2021). Inactivation Kinetics of Coxiella burnetii During High-Temperature Short-Time Pasteurization of Milk. Frontiers in microbiology, 12.

